# A conserved translation factor is required for optimal synthesis of a membrane protein family in mycobacteria

**DOI:** 10.1101/872341

**Authors:** Skye R.S. Fishbein, Ian D. Wolf, Charles L. Dulberger, Albert Wang, Hasmik Keshishian, Luke Wallace, Steven A. Carr, Thomas R. Ioerger, E. Hesper Rego, Eric J. Rubin

## Abstract

Ribosomes require the activity of associated GTPases to synthesize proteins. Despite strong evolutionary conservation, the roles of many of these remain unknown. For example, LepA (also known as elongation factor 4) is a ribosome-associated GTPase found in bacteria, mitochondria, and chloroplasts, yet its physiological contribution to cell survival is not clear. Recently, we found that loss of *lepA* in *Mycobacterium smegmatis* (Msm) altered tolerance to rifampin, a drug that targets a non-ribosomal cellular process. To uncover the determinants of LepA-mediated drug tolerance, we characterized the whole-cell proteomes and transcriptomes of a *lepA* deletion mutant relative to a wild-type strain. We find that LepA is important for the steady-state abundance of an outer membrane porin, which is integral to nutrient uptake and drug susceptibility. Loss of LepA leads to a decreased amount of porin in the membrane, resulting in the drug tolerance phenotype of the *lepA* mutant. LepA control requires a sequence motif in the 5’ region of the porin transcript. Thus, LepA controls the abundance of specific proteins, likely through its activity during translation.

**Importance:** Our understanding of how ribosomes properly synthesis an entire cellular proteome, in all its complexity, is still evolving. Ribosomal GTPases are often highly conserved, but the roles of many are not well understood. For example, elongation factor 4, or LepA, is a ribosome-associated GTPase conserved across bacteria, mitochondria, and chloroplasts. Using whole-cell proteomics and RNA-sequencing of wild type and a lepA deletion mutant, we find that LepA improves translation of mycobacterial porins in a message-specific manner. As porins play a key role in cell wall permeability, loss of LepA produces a plethora of phenotypic changes. These findings underline the problem of building proteins into a complex cell wall, such as that of mycobacteria, and point to a solution in the use of GTPases such as LepA, that have evolved to aid in specific protein synthesis.

## Introduction

Bacterial cells rely on transcription and translation to create a homeostatic proteome that is suitable for cell growth, yet adaptable to changing environments. From our first understanding of the regulation of the *trp* operon [1], post-transcriptional responses were thought to provide additional input to adaptive, cellular responses. Indeed, the ribosome machinery and its associated factors/RNA species adapt the cell during periods of environmental transition [2] [3, 4]. Recent advances in techniques such as quantitative proteomics, ribosome profiling, and cryo-electron microscopy (cryo-EM) have revealed an intricate balance of mRNA sequences and associating proteins at the ribosome that are necessary for the homeostatic cellular proteome [5–7]. Regardless of growth environment, bacterial cells are likely composed of a heterogenous population of ribosomes [8, 9], with some inactive and others actively translating parts of the proteome [10, 11]. Associating factors and RNA species tune each ribosome, creating a cell with a spatially- and temporally-regulated proteome [8, 12].

The successful synthesis of a given protein at the ribosome is mediated both by sequence features that are intrinsic to the mRNA and ribosome-associated factors. Structural elements of the 5’ untranslated region (UTR) control the rate of initiation, both constitutively and conditionally [13–15]. Local codon bias, an indirect measure of mRNA structure and tRNA availability, can also control the folding and abundance of a protein [3, 16, 17]. However, the ribosome is not a static machine sensitive only to signals encoded in the mRNA. It is also subject to regulation through associating factors that govern the outcome of translation. These include initiation factors [18–21] and the elongation GTPases [22–24], many of which are necessary for the initiation and progression of peptide bond formation at the ribosome during translation. Elongation factor P (EF-P), BPI-inducible protein A (BipA), and elongation factor 4 (LepA) associate with the 70S ribosome during elongation. Their role in translation appears to be message- and, in some cases, organism-specific [25–27]. These accessory factors may control the quality and timing of protein synthesis, proving critical for cell survival [28–30].

LepA is a conserved ribosome-dependent GTPase that is present in cells of almost all organisms, from bacteria to human mitochondria [27, 31]. LepA uses four classical elongation factor protein domains to contact the ribosome and hydrolyze GTP. In fact, it occupies the same position on the 70S ribosome as elongation factor G (EF-G) [32]. The C-terminal domain of LepA makes contact with the A/P-site tRNA and, likely, alters positioning of the tRNA species on the ribosome [33–35]. While it is non-essential in most bacteria, in some species, LepA might confer some fitness benefit in certain growth conditions, such as altered cation concentrations or low pH [36–41]. Despite its conservation, the physiological role of LepA remains unclear. Two different functions have been proposed for LepA. In *E. coli*, loss of LepA results in decreased polysome formation as a consequence of altered initiation rate and ribosome assembly defects [34, 41, 42]. Alternatively, structural studies of multiple bacterial LepA homologs indicate that while it binds the same site as EF-G on the 70S ribosome, it may alter the conformation of the ribosome-tRNA complex [43–45] and, perhaps, participate in translational quality control.

One clue concerning LepA function comes from work that defined the genetic determinants of single cell heterogeneity and drug susceptibility in mycobacteria. Mutations in several genes altered the rate of antibiotic-mediated bacterial killing, including *lepA*. Loss-of-function mutations in *lepA* resulted in increased survival in the presence of rifampin as compared to wild type Msm [46].

Here, we investigate the mechanistic basis of altered drug susceptibility of the mycobacterial *lepA* mutant. We find that LepA augments the protein levels of members of the mycobacterial porin (Msp) family during translation. LepA activity is determined by elements of the 5’ mRNA sequence of porin transcripts. LepA deficiency results in decreased synthesis of MspA, the major porin in Msm, and a reduction in cell permeability as measured by dye accumulation and killing by certain antibiotics. Thus, we find that LepA acts as a message-specific translation factor in mycobacteria, and provide evidence for its role in membrane protein translation.

## Results

### Loss of LepA results in mycobacterial drug tolerance through its activity at the ribosome

In a previous screen, we found that strains carrying transposon insertions in *lepA* were predicted to be associated with decreased accumulation of a fluorescent dye, calcein AM, and decreased killing by rifampin, a first-line tuberculosis drug [46, 47]. To validate that loss of LepA was responsible for the observed phenotype, we constructed a deletion of *lepA* (*lepA*-), and a corresponding complemented strain in which we re-introduced *lepA* in singly copy at a phage integration site (*lepA*+).We found that the *lepA*-strain exhibited an approximately 2-fold decrease in calcein signal (Figure 1A). Loss of LepA also resulted in increased tolerance to rifampin and vancomycin (Figure 1B,C) [46], but had no effect on isoniazid or linezolid tolerance (Supplementary Figure 1A,B) and no effect on susceptibility to a variety of translation inhibitors (Supplementary Table 1).

**Figure 1.**
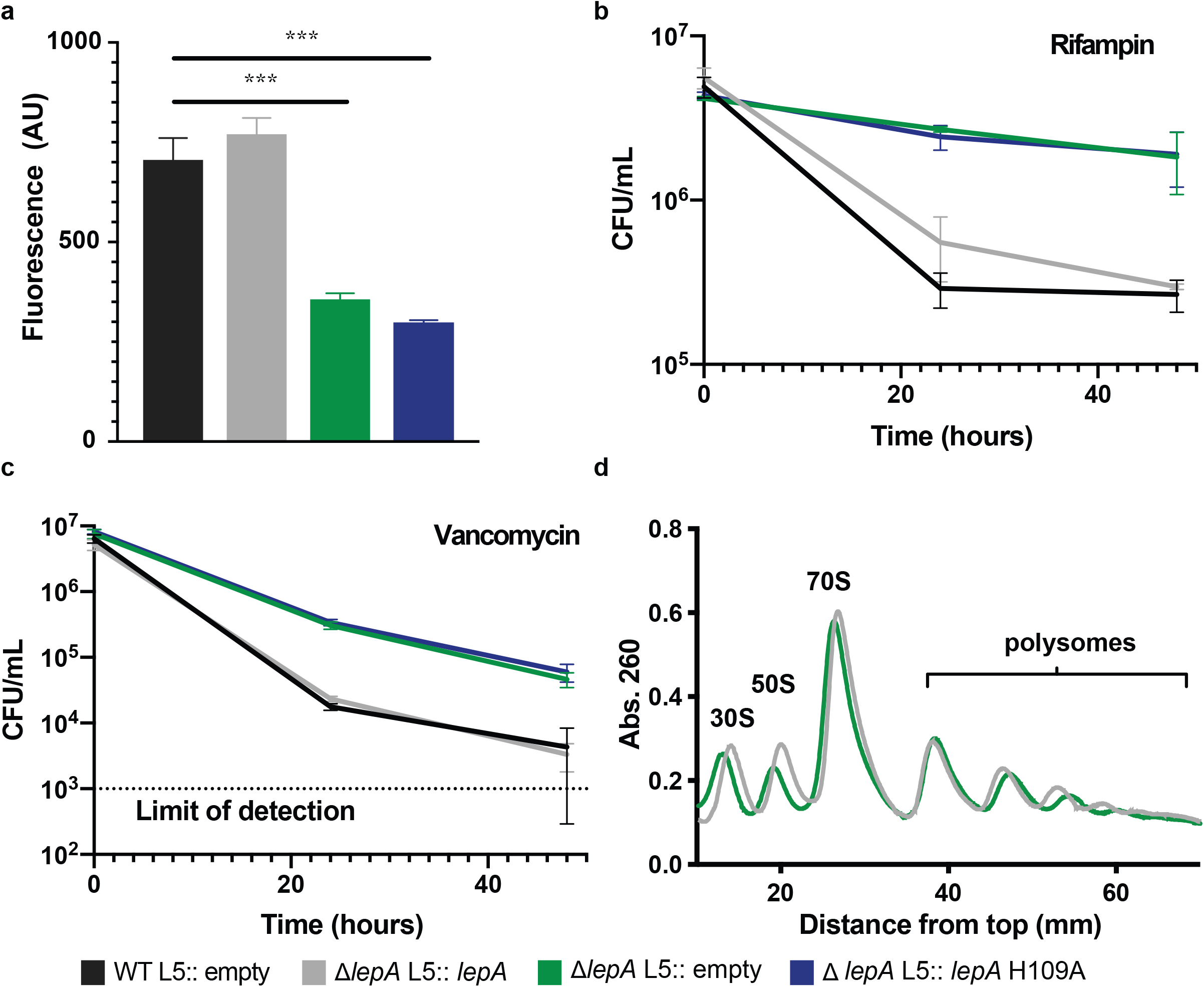
Loss of ribosome factor LepA causes altered drug tolerance in mycobacteria. a) Calcein staining across Msm strains with different *lepA* alleles. Values indicate mean calcein fluorescence across three replicates with error bars indicating standard deviation. ***P <0.001, calculated using a two-sided Student’s t-test. b-c) Msm *lepA* strains were treated with 10X MIC concentrations of rifampin and vancomycin, and cell survival was measured by colony forming units per mL (CFU/mL). All values are mean values with error bars indicating standard deviation of three biological replicates. d) Analysis of ribosome populations by sucrose density centrifugation and fractionation. Distance 0 corresponds to the lightest sucrose fraction.

To examine the link between a ribosomal factor and the phenotypes we observed, we first hypothesized that, like other LepA homologs [33, 41, 48], mycobacterial LepA functions as a ribosome-associated GTPase. To test whether the GTPase activity of LepA was critical to the phenotypes observed in *lepA*-cells, we assayed calcein staining and drug killing with a GTPase-null *lepA* allele (*lepA* H109A). Mutation of this conserved catalytic histidine was previously shown to abolish LepA GTPase activity in *E.coli* [42]. We found that both calcein levels and drug tolerance in the *lepA* H109A strain were equivalent to those in the *lepA*-strain (Figure 1A-C), indicating that, in mycobacteria, the effect of LepA on both calcein levels and drug killing was due to its GTPase activity.

We reasoned that LepA could be interacting with the ribosome during ribosome biogenesis, translation initiation, or elongation [37, 38, 42]. To determine if mycobacterial LepA was altering ribosome biogenesis, we examined ribosome populations in *lepA*+ and *lepA*-cells. Unlike in *E. coli*,loss of LepA in Msm did not alter levels of ribosome subunits or polysome formation, a marker of active translation (Figure 1D). To directly test if Msm LepA affects translation at the ribosome, we purified Msm LepA (Supplementary Figure 1C) and assessed its impact on the expression of a fluorescent protein (Venus) in a cell-free transcription-translation system. To test the influence of Msm LepA on translation, we added purified Venus mRNA to reactions with LepA at different concentrations. As previously observed with *E. coli* LepA [31, 49], addition of mycobacterial LepA to *in vitro* translation reactions increased the yield of fluorescent protein (Supplemental Figure 1D). Therefore, any effect on the yield of fluorescent protein must be due to the interaction of LepA with assembled ribosomes. These data suggest that in mycobacteria, LepA acts on active ribosomes during translation initiation or elongation, rather than during their biogenesis.

### Whole cell profiling of *lepA* mutant reveals dysregulation of outer membrane mycobacterial porins

Based on the altered drug tolerance of the *lepA* mutant, we hypothesized that LepA might affect the translation of proteins that mediate drug susceptibility. To find candidate proteins whose translation was affected by the loss of LepA, we measured simultaneous steady state levels of proteins and corresponding transcripts from wild type, *lepA*-, and *lepA*+ cells. To quantify the relative abundance of proteins, we used tandem-mass-tag (TMT) labeling of peptides, after tryptic digestion of cell lysates, coupled with LC-MS/MS. We identified a total of 4646 proteins with 4549 of them quantified by 2 or more peptides (Dataset 1). Among these 4549 proteins, 78 were significantly altered by the loss of LepA (Figure 2A). Interestingly, a number of membrane processes were enriched in the subset of proteins altered by LepA (Supplementary Figure 2A) [50]. Of these proteins, one of the most significant changes in the *lepA*-strain was a decreased abundance of peptides corresponding to a highly similar set of four porins: MspA, B, C, D, (Supplementary Figure 2B). Mycobacterial porins are octameric channels built into the outer membrane and are responsible for the uptake of nutrients critical for mycobacterial growth [51–55]. The four porins in Msm, encoded by *mspA-mspD*, are paralogs distributed across the genome. Each porin transcript is leadered, and encodes a Sec signal peptide which enables co-translational secretion of these proteins into the outer mycobacterial membrane [56, 57]. As the Msp porins are known to mediate drug susceptibility, likely through drug accumulation [58, 59], we hypothesized that LepA was affecting the abundance of these porins, contributing to the altered drug susceptibly we observed in the *lepA* mutant. Thus, we chose to concentrate our study on the LepA-mediated translation of the Msp porins.

**Figure 2.**
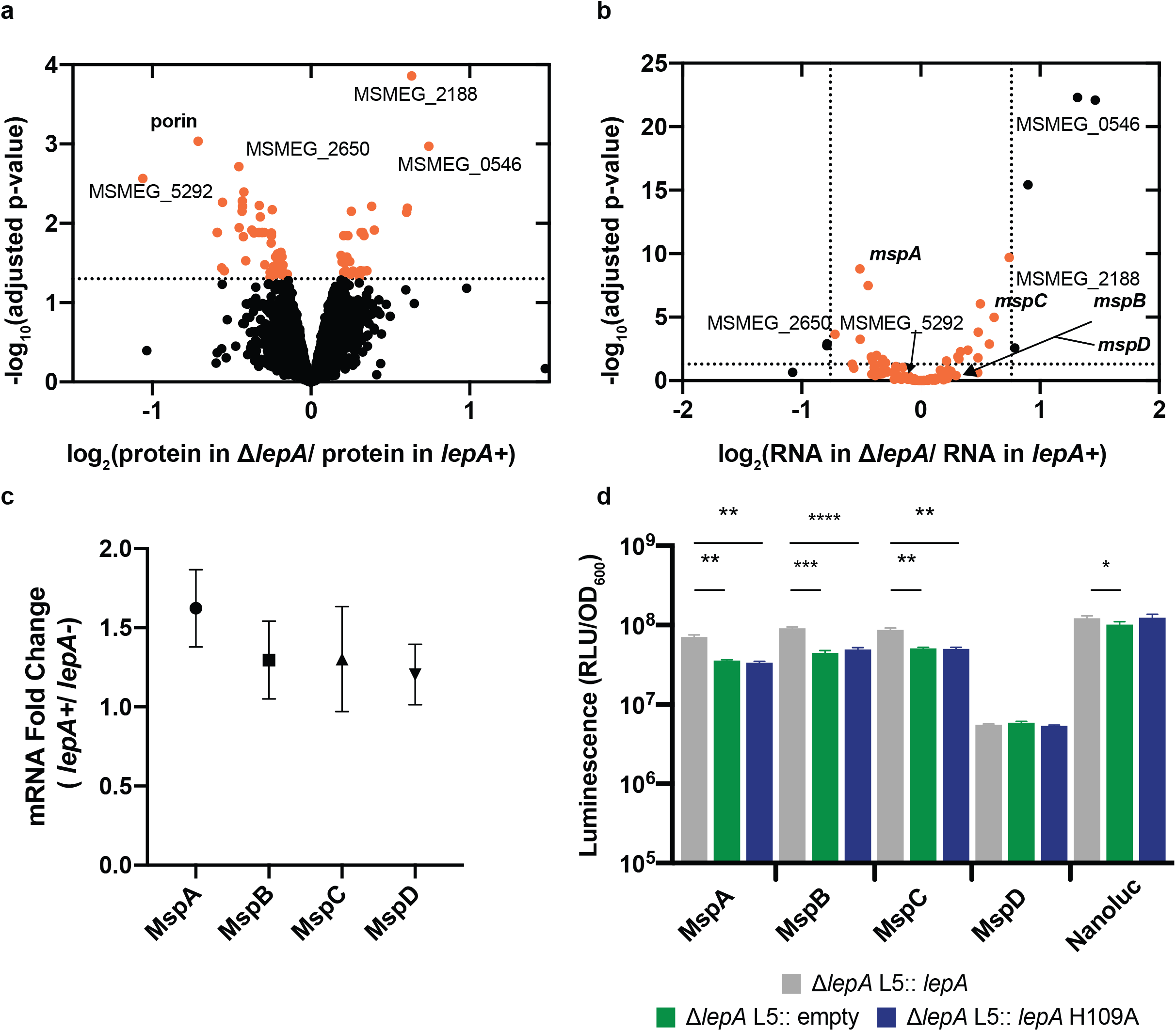
Whole cell profiling finds mycobacterial porins altered by loss of LepA. a) Log normalized ratios of protein represent reporter ion intensities in *lepA* knockout normalized by averaged reporter ion intensites from the strains containing *lepA* (Δ*lepA* L5:: *lepA* and WT L5:: empty). Orange dots indicate protein candidates that were significantly altered by loss of LepA. Bolded labels indicate genes/proteins of the mycobacterial porin family. ‘Porin’ indicates the collection of peptides that map to 4 proteins: MspA, MspB, MspC, and MspD. P-values for proteomic ratios were calculated using Student’s two-sided t-test, and adjusted for multiple testing using the Benjamini-Hochberg correction with an α of 0.05. b) Transcripts visualized correspond to protein messages significantly altered. Log normalized ratios of RNA represent RNA level *lepA* knockout normalized by averaged RNA level the strains containing *lepA* (Δ*lepA* L5:: *lepA* and WT L5:: empty). Orange dots represent transcripts with <1.7-fold change due to loss of LepA. Fold changes and adjusted p-values for RNA levels were generated using DEseq2.1.8. c) Changes in porin transcript levels due to LepA, as measured by RT-qPCR. Relative fold change of each mRNA was quantified by normalizing levels of each porin transcript to levels of *sigA* transcript. Points represent the mean of three biological replicates, with error bars depicting standard deviation. Fold changes were not significantly different from 1, as examined by a one-sided student t-test. d) Luminescence of porin reporters in Msm strains. Mean luminescence is depicted with error bars representing standard deviation of three biological replicates. ***P<0.001, ***P<0.01, *P<0.05, calculated using a one-way ANOVA, where each group was compared to the *lepA*+ strain. P-values were adjusted for multiple comparisons using a Bonferroni correction. Data in c,d are representative of multiple experiments.

To confirm that a change in the protein abundance was not caused by a change in transcript level, we used RNA-sequencing to estimate the transcriptional contribution to the set of regulated proteins from our proteomics (Dataset 2). While we found a number of mRNAs encoding membrane proteins were increased in *lepA*-strain, transcript levels across the four porin transcripts were altered in both directions, which was not consistent with our observation of the bulk decrease in protein levels of the porin family (Figure 2B). Indeed, to precisely measure the levels of each porin transcript, we used quantitative RT-PCR (RT-qPCR) and found that none were significantly altered by loss of LepA (Figure 2C). Together, these data show that loss of *lepA* leads to a disproportionate decrease in porin-derived peptides as compared to their encoding transcripts.

### LepA controls permeability by increasing MspA abundance in the membrane

As our whole-cell proteomics data identified peptides that mapped to all four porins, we sought to identify the porin whose translation was most affected by LepA. We fused each protein to a C-terminal luciferase reporter and expressed the fusions in single copy in a merodiploid, a strain that continues to produce the wild type copies of each protein. Using luminescence levels as a proxy for protein abundance, we examined levels of each porin in the absence of LepA or in the presence of a wild type copy or the GTPase null copy of LepA. The presence of functional LepA, but not the GTPase mutant (LepA H109A) increased luminescence 2-3 fold for fusions with coding sequences of MspA, MspB, and MspC, but not for MspD or luciferase alone (Figure 2D).

To further understand the cellular context of LepA-mediated control of Msp porins, we employed an inducible CRISPRi strategy to transcriptionally deplete each porin individually (Supplementary Figure 2C) [60]. To assess permeability, we compared calcein fluorescence of each porin knockdown in the presence or absence of LepA. Only depletion of *mspA* eliminated the LepA-dependent increase in calcein signal (Figure 3A). *mspA* is one of the most abundant transcripts in the cell and is the most highly expressed of the porins [61]. These data suggest that LepA-mediated regulation of MspA abundance is primarily responsible for the observed *lepA* deletion phenotypes. As altered permeability of *mspA* mutants can be assessed by a decrease in calcein and ethidium bromide fluorescence (Figure 3B,C)[62], we reasoned that if *mspA* and *lepA* were functioning in the same pathway, we should be able to detect this by an epistatic experiment. Accordingly, we deleted *mspA* in the original *lepA* deletion strain and measured fluorescence of both calcein and ethidium bromide (EtBr). We found that the loss of both genes does not lead to a more severe reduction in calcein accumulation that the single *mspA* mutant. Thus, *lepA* is epistatic to *mspA* in Msm, suggesting they function in the same pathway. Supporting this conclusion, we find that an MspA-mRFP fusion is less abundant at the membrane in the absence of LepA (Supplementary Figure 2D). Collectively, these data suggest that LepA is sufficient to affect the translation of multiple porins, but that it mainly controls the translation of MspA, which in turn mediates permeability to multiple compounds, including antibiotics.

**Figure 3.**
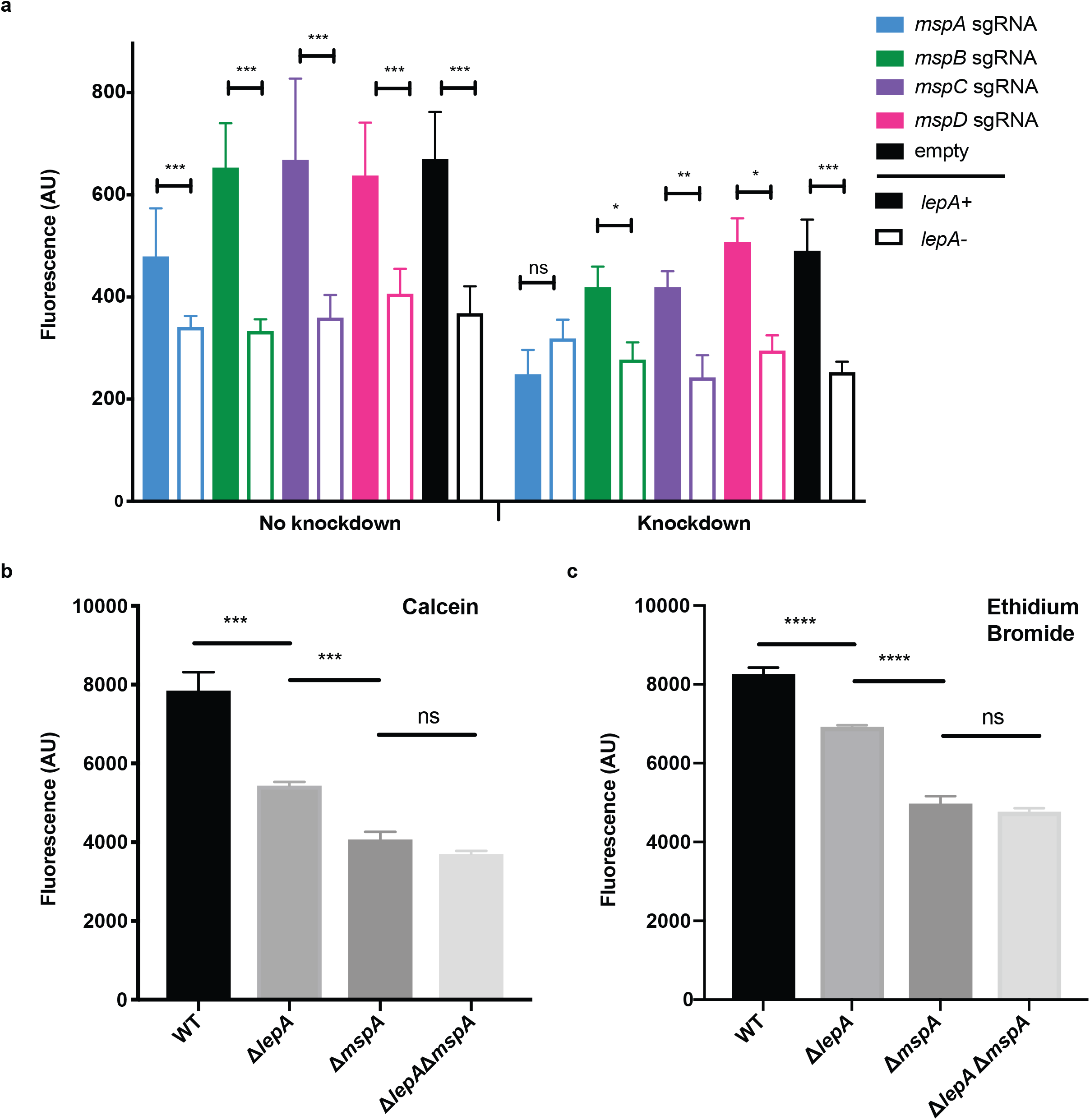
Loss of LepA causes membrane defects primarily through control of MspA. a) Calcein staining of *lepA*-porin genotypes. Knockdown of each porin was assessed in a *lepA*+ strain (filled bar) and in a *lepA*-strain (open bar). Mean fluorescence is depicted with error bars indicating standard deviation across three biological replicates. Sets of colors bars denote porin-specific knockdown strains. ‘Empty’ refers to strains containing the control vector with the aTc-inducible CRISPRi system and no target-specific sgRNA. ***P<0.001, **P<0.01, *P<0.05; calculated using a twosided Student’s t-test. b) Calcein fluorescence across Msm strains, with error bars indicating standard deviation across three biological replicates. c) EtBr fluorescence across Msm strains, with error bars indiciating standard deviation across three biological replicates. ***P<0.001, **P<0.01, *P<0.05, calculated using a one-way ANOVA, with p-values adjusted for multiple testing correction using Bonferroni correction.

### LepA affects MspA abundance through mRNA sequence determinants

How does LepA recognize MspA as a target? LepA has previously been shown to interact with the peptidyl-transferase center (PTC) of the 70S ribosome, and increase the amount of protein synthesized [33, 63]. Because our data suggest that Msm LepA functions during translation to selectively increase the abundance of a subset of porins, we hypothesized that the LepA-controlled porins have sequence features that allow for LepA control of translation rate. Supporting this notion, we find that in the absence of LepA, *mspA* transcript was less stable, likely because of less efficient translation leading to increased transcript degradation [64] (Figure 4A). To define LepA-related features of the porin, within either the mRNA sequence or the protein sequence, we took advantage of the fact that LepA affects the abundance of MspA-C and not MspD. Previous LepA ribosome profiling data from *E.coli* implicated the presence of certain glycine codons in transcripts with an altered translation rate [41, 42]. We found a 2-3 fold increase in the use of the glycine codon ‘GGT’ in *mspA-C* relative to *mspD* (Supplementary Figure 3A). However, we found that replacing these with synonymous codons did not abolish the LepA-dependent increase in luminescence signal (Supplementary Figure 3B,C). Although the amino acid sequences of MspA and MspD have a high average pairwise-identity (82%), the nucleotide sequences are less similar (average pairwise identity of 75%). Therefore, we hypothesized that variants in the transcript sequence may contribute to LepA-dependence. We randomly re-assigned codons in the *mspA* gene by sampling from two different probability distributions: the codon bias of the Msm genome (GC-rich) and the inverse codon bias relative to the Msm genome (AT-rich). Changing the transcript sequence decreased the effect of LepA on porin reporter abundance (Figure 4B). Thus, LepA modulation of MspA abundance occurs through sequence determinants in the mRNA.

**Figure 4.**
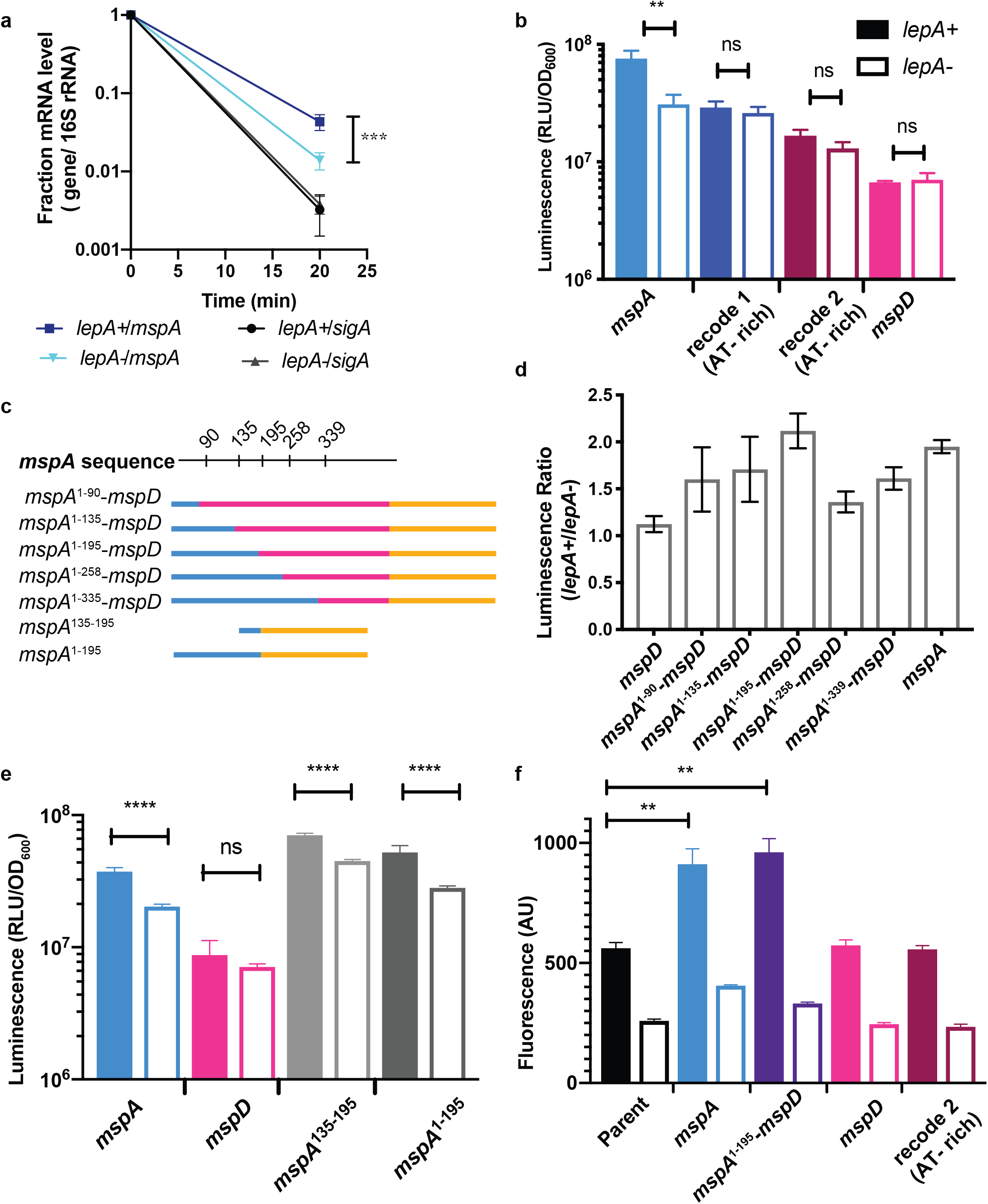
LepA affects permeability through mRNA-sequence determinants in *mspA*. a) Transcription was inhibited at time 0 (t=0) to examine *mspA* and *sigA* stability over time, by RT-qPCR. Fraction mRNA level indicates the relative level of a specific mRNA (*mspA* or *sigA*) normalized to its level at t=0, with each point representing average and error bars indicating standard deviation around three replicates. For each transcript, endpoint fractional abundances in *lepA*+ were compared to those in *lepA*-using a student t-test. ***P<0.001. Data are representative of multiple experiments. b) Luminescence of reporters where *mspA* was recoded using a Msm codon bias (GC-rich recode), and a less optimized codon bias (AT-rich recode). Bars indicate mean luminescence with error bars indicating standard deviation around three replicates. Data are representative of multiple experiments. c) Schematic for porin chimeras used in experiments. Blue parts of the gene represent regions of the protein encoded by *mspA* sequences, and pink parts of the gene represent regions of the protein encoded by *mspD* sequences. d) Luminescence was measured for porin fusions in (c). Bars indicate ratio of mean luminescence for each porin chimera, with error bars indicating standard error about the mean. Means represent average of three biological replicates. e) Luminescence was measured for chimeric/fragment porin sequences. Bars indicate mean luminescence with error bars indicating standard deviation around three replicates. Data are representative of multiple experiments. f) Calcein staining of luciferase-porin reporter strains from b-d) was used to assess porin coding sequence contribution to permeability. ‘Parent’ indicates the absence of any additional reporter, thus *lepA*+ (filled bar) and *lepA*-strains (open bar). Each subsequent set of bars indicates derivatives of those two parental strains containing the specified porin reporter. Bars indicate mean fluorescence with error bars indicating standard deviation around three replicates. ****P<0.0001, **P<0.01 calculated using a one-sided ANOVA.

To determine the location of such sequences that may confer LepA control to a coding sequence, we compared MspA-D sequences in search of regions that might determine LepA control. The alignment of MspA-D (Supplementary Figure 2B) suggests that the most divergent part of the sequence resides in the signal sequence region (the first 30 codons). To test whether a LepA-mediated increase in porin abundance is due to the nature of the signal sequence, predicted signal peptide coding sequences were translationally fused to the luciferase gene, and expressed in the presence or absence of LepA. We found that the presence or absence of LepA did not affect production of MspA-D signal sequences through luminescence (Supplementary Figure 3D). To further investigate sequence determinants, we constructed chimeras of *mspA* and *mspD* to map the location in the coding region that was LepA-sensitive. Each chimera contains *mspA* coding sequence at the 5’ end and *mspD* coding sequence at the 3’ end (Supplementary Figure 3E, Figure 4C), followed by a luciferase fusion. We found the strongest influence of LepA on *mspA*^1-195^-*mspD*, with *mspA*^1-135^-*mspD* also displaying some LepA-mediated increase in luminescence (Figure 4D). This indicated that a sequence determinant within the first 195bp, but after the signal sequence, was responsible for the LepA-dependent increase in expression. Additionally, we noted the decrease in LepA-dependent signal in the *mspA*^1-258^-*mspD* chimera, indicating that the sequence determinants required for optimal LepA effect are complex.

To further define the contribution of this ~100bp region to LepA control of reporter signal, we constructed a series of chimeras containing *mspA* sequence fragments followed by the luciferase reporter (Figure 4C). The fusion containing the first 195 bp of *mspA* alone produced a statistically significant increase in protein due to LepA that was biologically comparable to the increase in MspA levels due to LepA (Figure 4E). Taken together, these data suggest that the LepA control of MspA abundance requires an mRNA sequence near the 5’ end of the transcript, immediately downstream of the signal sequence. We hypothesized that the presence of this sequence-based control contributed to the synthesis of a functional porin in Msm. To examine the influence of mRNA elements on cellular permeability through LepA, we used the previously-constructed luciferase fusion reporters, all expressed from the same strong constitutive promoter. At baseline, the *lepA*+ strains displayed higher calcein fluorescence than those that lacked *lepA*. Yet, the addition of an extra porin did not always increase calcein fluorescence correspondingly. Only the presence of the mRNA elements, which increase reporter levels through LepA, correlated with an increase in calcein staining (Figure 4F). Specifically, the addition of LepA-controlled porin reporters, such as MspA-luciferase or MspA^1-195^-MspD-luciferase, increased calcein permeability. Porin reporters, whose levels were unaffected by LepA, such as the recoded MspA-luciferase and MspD-luciferase, did not increase the permeability of the mycobacterial cells, and the presence of LepA did not augment calcein permeability in these strains. These data suggest that production of a functional porin in the outer membrane depends on the both the activity of LepA and corresponding codon bias in the porin transcript.

## Discussion

LepA is highly conserved across the kingdoms of life, yet its cellular role remains unclear. In *E. coli*, LepA globally influences translation through ribosome biogenesis, something we did not observe in Msm [34, 42]. Instead we find that in Msm, LepA influences the abundance of porins, the major outer membrane proteins of this organism. While the *lepA* mutants are viable, they have a very specific permeability defect. This finding suggests a rather specialized function for LepA. We observe transcriptional upregulation of a number of membrane/transport proteins in the absence of LepA, further underlining the membrane defect as the major consequence of losing LepA. Additionally, a number of observations exist from genome-wide transposon screens, indicating that LepA plays a critical role, likely at the membrane. In fact, these data suggest, that while *lepA* is non-essential for *in vitro* growth of *Mycobacterium tuberculosis* (Mtb), *lepA*-strains grow poorly during murine infection [65]. We speculate that synthesis of membrane proteins may be increased or is more costly during infection, necessitating the function of LepA. Further, in Mtb, *lepA* becomes essential for growth in the absence of *ponA1*, a prominent cell-wall biosynthetic enzyme in Mtb [66]. Loss of such cell-wall enzymes may result in collateral perturbation to the cell membrane proteome, causing LepA to become indispensable to the cell. LepA function in mycobacterial appears to be critical for maintenance of the mycomembrane, a very complex network of lipids and proteins [67].

How does LepA control the abundance of MspA? We find that LepA alters the abundance of specific porins independent of the promoter and UTR elements. One model is that LepA affects elongation along mRNA messages. Specifically, we observed that approximately 100bp of the 5’ end of the open reading frame (ORF) was required for LepA-mediated modulation. The 5’ region of a message may encode a number of signals that dictate the level of protein made from a given message, including mRNA structure and codon bias [68–71]. Synonymous recoding of *mspA* across the gene abolished LepA-dependent reporter expression, suggesting that the information in the mRNA sequence is necessary for LepA activity on a porin message. Non-optimal codons, for example, are posited to alter ribosome speed at beginning of translation to accommodate loading/assembly of multiple ribosomes onto a message [72, 73]. LepA’s role as a translational GTPase may be selected for regions of transcripts made up of certain codons.

Altering the translation rate could have profound effects on protein folding, particularly for a secreted protein such as a porin. Sequence elements that control translation rate can help accommodate the initiation of Sec-dependent translation as the nascent peptide emerges from the exit tunnel [74–76]. Binding of LepA to the ribosome, facilitated by a series of rare codons, could alter the rate of translation, thus changing the folding of the nascent peptide and affecting its ability to be recognized by the secretory system or other factors (Figure 5). Certainly, a ribosome carrying a nascent signal peptide can interact with a number of distinct components of the Sec system, which enable co-translational secretion of membrane-destined proteins [10, 77, 78]. It is well-established that translation rate is intimately tied to protein folding [79], thus, we do not divorce LepA’s activity from either of these outcomes. We suspect that the LepA’s interaction with the *mspA* message results in a change in translation rate, affecting the folding of the nascent, secreted peptide into the membrane.

**Figure 5.**
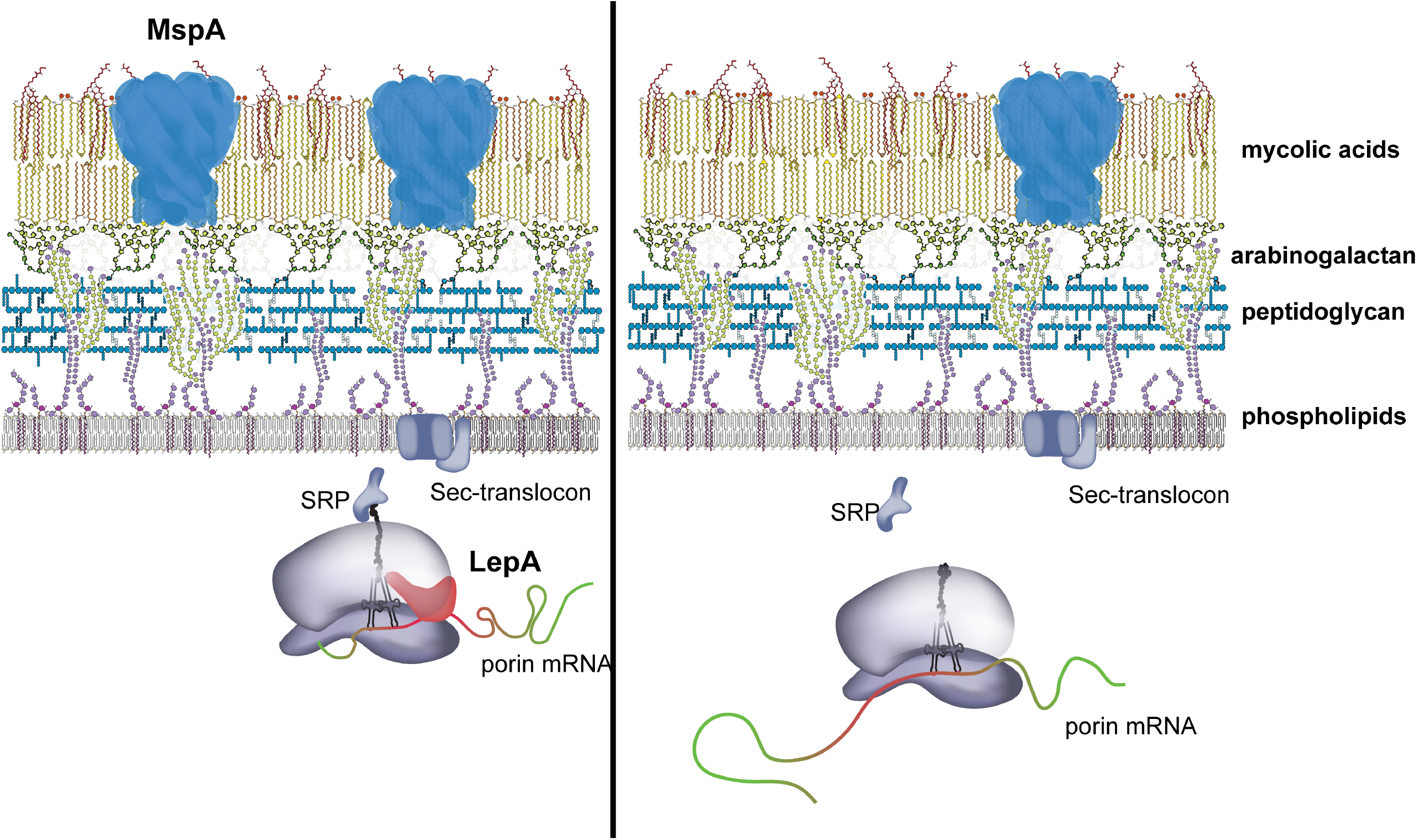
Mechanism of LepA control of permeability. Model of LepA modulation of porin translation. LepA aids translation of outer membrane porins, most notably MspA, based on a series of codons in the 5’ region of *mspA* transcripts. LepA-determining mRNA sequences of the porin transcript are in red, while the rest of the transcript is in green. SRP, signal recognition particle

What then is the physiologic role of LepA and the functional significance controlling protein synthesis for a subset of proteins? Our data suggest that LepA has a constitutive role in aiding the translation and folding of a set of proteins with particular characteristics. This model implies that some proteins are either difficult to make, fold or insert into their proper niche in the cell, e.g., the outer membrane. MspA is one of the most abundant transcripts in the cell; therefore, it is plausible that a significant proportion of ribosomes are translating MspA at any given time. Improper synthesis of MspA, either through the toxic protein product, or stress on ribosome populations, could represent a significant stress on the cell and justify a role for a more or less dedicated factor to aid in translation and folding.

Why is LepA so conserved? In other cellular systems, LepA’s function appears to be related to expression of abundant membrane proteins. In mitochondria, LepA is associated with the membrane and alters the levels of inner membrane respiration complexes through its activity at the ribosome [37, 48, 80]. In *Arabidopsis thaliana* chloroplasts, LepA associates to the thylakoid membrane, and its absence was associated with a reduced production of photosystems I and II components [36]. It appears that LepA plays a similar role in these different organisms and organelles, contributing to the production of important membrane structures. In bacteria, the location of sequence determinants for LepA suggests that LepA may play a role in helping to coordinate translation with signal peptide recognition and trafficking of the ribosome to the membrane. While organelles such as the mitochondria do not possess the SRP pathway, these organelles have systems that play similar roles integrating proteins into membranes directly from the ribosome [81, 82]. Furthermore, porins, respiration complexes, and photosystems are all abundant, multi-unit complexes of proteins, which are all known to be post-transcriptionally controlled to ensure proper localization, folding, and levels of the complexes [83, 84]. Thus, LepA might serve as a quality-control measure that operates across phylogeny to ensure that even the most abundant and difficult proteins are produced efficiently.

## Acknowledgements

We thank Dr. Allison Carey and Dr. Thibault Barbier for advice on the manuscript. We also thank Dr. Allen Buskirk for his reading of the manuscript. We thank Dr. Sarah Fortune and the Fortune lab for insightful discussion on the work. We thank Dr. Bill Neidermeyer, and members of the Whelan lab for allowing us to use ribosome fractionation machinery. We are grateful to the Dr. Michael Welsh and the Walker lab for allowing us to use their protein purification equipment. This research was supported by funding from the NIH (U19 AI107774 to E.J.R.).

## Author Contributions

Conceptualization: S.R.S.F, E.H.R, C.L.D, E.J.R; Methodology: S.R.S.F, E.H.R, C.L.D, E.J.R, H.K., T.R.I, S.A.C.; Investigation: S.R.S.F, E.H.R, I.D.W, A.W, L.W.,H.K.,T.R.I,; Data Curation: S.R.S.F, H.K.,T.R.I; Writing – Original Draft: S.R.S.F, E.H.R, E.J.R.; Writing – Review & Editing: S.R.S.F, E.H.R, E.J.R., H.K., T.R.I., S.A.C., C.L.D, A.C., and T.B.

## Materials and Methods

### Bacterial strains and growth conditions

Msm strains were inoculated from frozen stocks into Middlebrook 7H9 medium supplemented with 0.2% glycerol, 0.05% Tween-80, and ADC (5g/L bovine serum albumin, 2g/L dextrose, 3 mg/L catalase) and grown at 37°C. Appropriate antibiotics or inducing agents were used at the following concentrations in Msm: nourseothricin (Nat, 20 μg/ml), zeocin (Zeo, 20 μg/ml), kanamycin (Kan, 25 μg/ml), hygromycin B (Hyg, 50 μg/ml), anhydrous tetracycline (aTc, 100 ng/ml). Transformations, performed for the construction of Msm strains, were plated onto LB (Luria Broth) (LB) agarose plates supplemented with the appropriate antibiotic. Unless otherwise specified for an experiment, strains were grown to log-phase (OD_600_ 0.3-0.8) without antibiotics. For cloning purposes, *E. coli* strains were grown in LB broth or on LB agar with antibiotics used at the following concentrations: Nat (40 μg/ml), Zeo (50μg/ml), Kan (25 μg/ml), and Hyg (100 μg/ml).

### Bacterial strain construction

All bacterial strains constructed in this study are found in Supplementary Table 2. Description of the plasmids, primers, and recombinant DNA used to construct the strains can be found in Supplementary Table 3, 4, and 5 (respectively). Generally, all plasmids were constructed by restriction digestion of the parental vector (with the desired antibiotic resistance gene and phage integration gene for Msm propagation) and all inserts were prepared by amplifying gene fragments with 18-25bp-Gibson assembly overhangs. Vector and insert combinations were fused together by Gibson isothermal assembly [85]. Plasmids were isolated from *E. coli* and insert orientation was sequenced via Sanger sequencing. Sequencing reactions were carried out with an ABI3730xl DNA analyzer at the DNA Resource Core of Dana-Farber/Harvard Cancer Center (funded in part by NCI Cancer Center support grant 2P30CA006516-48).

#### Deletion mutants

The *lepA* mutant, Δ*lepA*::zeo (HR334), and the *mspA* mutant, Δ*mspA*::zeo (HR329) were a generous gift from E.H. Rego [46]. The Δ*lepA*::zeo Δ*mspA*::hyg strain (SF789) was constructed with HR334 as the parental strain, using double-stranded recombineering. For recombination, a linear dsDNA fragment was generated by amplifying the following fragments: 500 bp upstream region of *mspA*, 500 bp downstream region of *mspA*, a *lox*-hyg-*lox* fragment. The three fragments were ligated together using Gibson assembly. The deletion cassette was transformed in to an Msm recombineering strain as previously described [86, 87], and plated on Hyg to select for double mutants.

#### *lepA* and *mspA* alleles

Plasmid pSF121, used for *lepA* complementation was generated using a parental vector (pCT94) that integrates into the L5-phage site, and is marked with a Kan-resistance (kan) gene. The vector was digested with XbaI and HindIII (NEB). *lepA* and its 5’ UTR (300bp upstream) were amplified via PCR. The vector and insert were gel extracted and ligated using Gibson assembly. The complemented strain, *lepA+* (SF178: Δ*lepA*::zeo, L5::*lepA*-kan) and marked mutant, *lepA*- (SF181: Δ*lepA*::zeo, L5::*empty*-kan) were derivatives of HR334, respectively. Plasmid pSF417, used for *lepA* complementation in the CRISPRi experiments, was generated using a parental vector (pCT204) that integrates into the Tweety(Tw)-phage integration site and is marked with a Hyg-resistance (hyg) gene. The *lepA* insert was amplified from the Msm genome as above, and the parental vector was linearized by digesting with SspI and NdeI. The resulting vector was assembled as above.

For microscopy, *mspA* was fused to mRFP, with no linker, constitutively expressed from P_rpsA_, in a Tw - integrating vector with a nourseothricin-resistance (nat) gene. The *mspA*-mRFP vector was transformed into *lepA*+ and *lepA*-for localization of MspA-mRFP.

#### Candidate-luciferase fusions

All luciferase reporters were generated from the parental vector CT250, a Tw-integrating vector. The vector was linearized using NdeI and HindIII, to preserve the upstream promoter (P_rpsA_). For each reporter, the candidate gene was amplified from the Msm genome and the luciferase gene was amplified from a plasmid (pJR976) kindly provided by Jeremy Rock [60]. Using Gibson overhangs that contained glycine-serine-glycine (GSG) linkers, each reporter vector was constructed using Gibson assembly. Each reporter vector was transformed into SF178 or SF181.

#### Porin-knockdown constructs

Knockdown of each porin was accomplished using mycobacterial CRISPRi, with knockdown systems constructed as described [60]. Porin knockdown vectors were created by annealing oligos for each porin and ligating these fragments into a linearized vector (pJR965 digested by BsmBI), containing the mycobacterial CRISPRi system. Knockdown-vectors were co-transformed into HR334 with pSF417 or pSF418.

#### Signal sequence fusions

The SignalP server [88] was used to predict the length of signal peptides in MspA-D. Signal sequences from each porin (the first 84bp from *mspA*, 87bp from *mspB*, 96bp from *mspC*, and 75bp from *mspD*) were amplified from the Msm genome and cloned in-frame upstream of the GSG-luciferase reporter gene as previously described, to create pSF413, pSF414, pSF415, and pSF416, respectively.

#### Chimeric fusions

In-frame fusions of *mspA* and *mspD* were built to examine sequence contribution to reporter signal. Each fusion point was selected, such that the codon on either side only contained synonymous differences between either porin sequence. For a given fusion point, denoted by arrows in Supplementary Figure 3E, the 5’ sequence of the chimera was amplified from *mspA* and the 3’ sequence of the chimera was amplified from *mspD*. As with other reporter experiments, the chimeric sequence was cloned 5’ to a GSG-luciferase reporter. Fragments for each chimera were annealed to a linearized CT250 (using NdeI and HindIII) using Gibson assembly.

### Calcein-AM staining

Strains were grown to log-phase and stained with 0.5 μg/ml of calcein-AM (Invitrogen, Carlsbad,CA) for one hour. Strains were analyzed by flow cytometry on a MACSQuant (VYB Excitation: 488nm; Emission filter: 525/50) in the same manner as previously described [46]. Median fluorescence was used from each replicate to compute an overall mean fluorescence intensity.

### Kill curves

Strains were grown to mid-log phase, diluted to OD_600_ 0.05 and treated with 10X MIC concentrations of the following drugs: rifampin (20 μg/ml), isoniazid (40 μg/ml), vancomycin (4 μg/ml) and linezolid (500 ng/ml). Survival was assessed over time as described previously [46].

### Drug susceptibility assays

Drug susceptibility was determined using a minimum inhibitory concentration (MIC) assay, as described previously [89]. In 96-well plates, strains were diluted to 0.005 and tested in biological triplicate in serial dilutions of tetracycline (Sigma Aldrich, St. Louis, MO), clarithromycin (Sigma Aldrich, St. Louis, MO), chloramphenicol (Sigma Aldrich, St. Louis, MO), amikacin (Sigma Aldrich, St. Louis, MO), and erythromycin (Santa Cruz Biotechnology, Santa Cruz, CA). The highest concentrations of each drug test were: 4 μg/ml for tetracycline, 4 μg/ml for clarithromycin, 320 μg/ml for chloramphenicol, 3.2 μg/ml for amikacin, and 16 μg/ml erythromycin. Plates were agitated at 37°C for 21 hours. To determine MICs for each condition, 0.0002% resazurin was added to each well and, plates were agitated at 37°C for 3 hours. The first well with no growth (blue) in each concentration gradient was considered the MIC. A biological replicate, in this case, is considered a single row, in a 96-well plate, of drug and bacterial incubation, using bacteria from the same culture.

### Purification of mycobacterial LepA

Msm LepA was cloned with an N-terminal 6x His-tag, using Gibson assembly and expressed from pET28a in BL21 *E. coli* as previously described for *E. coli* LepA [42]. Briefly, 1 L of log-phase culture was induced with 1 mM Isopropyl β-D-1-thiogalactopyranoside (IPTG) for 4 hours at room temperature. Cells were harvested at 5,000 g x 10 min, and pellets were frozen at −80°C overnight. The pellet was thawed with a stir bar in 30 mL of lysis buffer (50 mM Tris-HCl pH 7.6, 60 mM KCl, 5% glycerol, 6 mM β-mercaptoethanol (BME), 2 mM MgCl_2_, 0.2 mM phenylmethylsulfonyl fluoride (PMSF), DNase), and lysed by French press. Cell lysates were clarified by centrifugation at 15,000 x g for 30 min at 4°C. Lysate was brought up to 30 mM imidazole pH 7.6 and his-tagged LepA was extracted via batch binding with 4 mL Ni-NTA beads for 2 hours with a stir bar at 4°C. Beads were collected in plastic columns with a 10 mL bed volume and washed with 4x 10 mL wash buffer (50 mM Tris-HCl pH 7.6, 300 mM KCl, 5% glycerol, 6 mM BME, 30 mM imidazole), corresponding to wash 1 – 4.1 mL elution fractions were collected using elution buffer (50 mM Tris-HCl pH 7.6, 40 mM KCl, 5% glycerol, 6 mM BME, 200 mM imidazole) and analyzed via SDS-PAGE (Supplementary Figure 1C). The cleanest elution fractions (2, 6, and 7) were pooled and dialyzed into 6 L (3 x 2 L) of storage buffer (50 mM Tris-HCl pH 7.6, 50 mM KCl, 5% glycerol, 6mM BME) using dialysis cassettes with 10 kDa MWCO. 10 μL aliquots were flash frozen with liquid nitrogen and stored at −80°C. LepA protein concentration was calculated using the Bradford protein assay.

### *In vitro* translation

To assess the effect of LepA on translation, *in vitro* translation reactions were prepared with purified mRNA. Plasmid pSF741 was used in a HiScribe T7 *in vitro* transcription kit (New England Biolabs, Ipswich, MA) to generate Venus mRNA. A master mix of purified Venus mRNA (500 ng per reaction) and PureExpress (New England Biolabs, Ipswich, MA) components were prepared in duplicate reactions with a range of concentrations of purified Msm LepA. When no LepA was added to the reaction, an equal volume of storage buffer was added in place of protein. Reactions were carried out in 12.5 μL in a 384-well plate for 4 hours at 37°C, and fluorescence (measured at an excitation of 505 nm and an emission of 540 nm) was collected on a TECAN Spark 10M plate reader.

### Ribosome analysis

Preparation of mycobacterial ribosomes was performed as previously described for *E. coli* [90], yet optimized for Msm. 500 ml of cells were grown to mid-log phase, filtered over 0.22 μm, 90 mm membranes (Millipore) on a fritted glass microfiltration apparatus (Kimball-Chase), and scraped into liquid nitrogen. 500 μL of lysis buffer (20mM Tris pH 8, 10 mM MgCl_2_, 100 mM NH_4_Cl, 5mM CaCl_2_, 0.4% Triton-X 100, 0.1% NP-40, 34 mg/ml chloramphenicol, 100 U/ml RNase-free DNase I) was added to cell scrapes. Frozen cells and lysis buffer were ground in a Retsch 400 mixer mill using 10 mL grinding jars and 12 mm grinding balls at 15 Hz for 5 x 3 minutes. Cell lysates were thawed and clarified at 15,000 x g for 15 minutes at 4°C. 250 μL of lysate were layered onto a 10-40% linear sucrose gradient. The sucrose gradients were spun in a Beckman ultracentrifuge at 35,000 rpm (150,000xg) for 2.5 hr at 4°C. The gradients were fractionated and analyzed using a gradient fractionator (BioComp Instruments, Inc., NB, Canada).

### Proteomics and RNA sequencing

60 mLs of each strain were grown to log-phase (OD_600_ ~ 0.4) and the culture was split into two parts to extract protein and RNA separately. Both aliquots were spun at 4000 rpm for 10 min. For proteomics, cells were resuspended in 500 μL of urea lysis buffer (8 M urea in 50 mM Tris pH 8.2, 75 mM NaCl, Roche Complete EDTA-free Protease Inhibitor Cocktail tablet) and subjected to bead-beating for 4 x 45 sec with 3 minutes on ice in between. Cell lysates were spun down at 10 min, 13,000 rpm at 4°C and the supernatant was isolated for proteomics sample preparation. For RNA sequencing, RNA was isolated as described previously [91], depleted for rRNA using RiboZero (Epicenter), and prepared for sequencing using KAPA Stranded RNA-Seq Library Preparation Kit (KAPA Biosystems).

#### Quantitative Proteomics

Samples for quantitative proteomics experiments were processed as described previously [92]. Briefly, three biological replicates of each strain’s lysates were reduced with 5 mM dithiothreitol (DTT), alkylated with 10 mM iodoacetamide (IAA), digested with Endoproteinase Lys-C (Wako Laboratories) for 2 hours at 1:50 enzyme to substrate ratio at 30°C, followed by an overnight digestion with trypsin (Promega) at 1:50 enzyme to substrate ratio at 37°C. Reactions were quenched with neat formic acid (FA) for a final concentration of 1%. Digests were desalted using tC18 SepPak reversed phase cartridges (Waters, Milford, MA) following manufacturer’s protocol. A tandem mass tag (TMT) isobaric labeling strategy was used for this experiment. 50ug aliquot of each of the 9 samples were labeled by TMT10plex reagent (ThermoFisher Scientific, Waltham, MA) following the manufacturer’s protocol. A pooled reference standard was generated by mixing equal amounts of each of the nine samples and included in the tenth channel of the TMT10plex. Labeling efficiency was assessed prior to quenching the reactions. Once sufficient (>99%) labeling efficiency was achieved, reactions were quenched and samples were mixed together. Combined sample was desalted using tC18 Sep-Pac reversed phase cartridges, and the eluate was dried down completely. Sample was reconstituted and fractionated on Zorbax 300 Extend-C18 4.6 x 250mm column (Agilent Technologies, Santa Clara, CA) as described previously [92]. Fractions were collected every minute during the gradient and further concatenated into a total of 24 fractions that were analyzed on Q Exactive Plus mass spectrometer (MS) coupled to EASY-nLC 1200 ultra-high performance liquid chromatography (UHPLC) system (ThermoFisher Scientific, Waltham, MA). One microgram of each of the fractions was injected on a 75um ID Picofrit column (New Objective, Woburn, MA) packed with Reprosil-Pur C18-AQ 1.9um beads (Dr. Maisch, GmbH) in-house to a length of 22cm. Sample was eluted at 200nL/min flow rate with solvent A of 0.1% FA /3% acetonitrile (ACN), solvent B of 0.1% FA / 90% ACN and a gradient of 2-6% B in 1min, 6-30% B in 84min, 30-60% B in 9min, 60-90% B in 1min, and a hold at 90% B for 5min. MS data was acquired in data-dependent mode with MS1 resolution of 70,000 and automatic gain control (AGC) of 3e6. MS/MS was performed on most intense 12 ions with a resolution of 35,000, AGC of 5e4, isolation width of 1.6amu, and normalized collision energy of 29. Data was extracted and searched against a *M. smegmatis* database using Spectrum Mill MS Proteomics Workbench (Agilent Technologies, Santa Clara, CA). Extracted spectra were searched using carbamidomethylation of cysteins and TMT labeling of N-termini and lysine residues as fixed modifications and methionine oxidation, asparagine deamidation and protein N-terminal acetylation as variable modifications. Spectrum to database matching was controlled with peptide level false discovery rate (FDR) of less than 1%. Peptides were rolled into protein groups and subgroups in Spectrum Mill with a protein level FDR of 0%. Protein summary export consisting of list of quantified proteins with reporter ion ratio of every TMT channel to the pooled reference channel was generated for quantitation of proteins. TMT10 reporter ion intensities were corrected for isotopic impurities in the Spectrum Mill protein/peptide summary module using the afRICA correction method which implements determinant calculations according to Cramer’s Rule and correction factors obtained from the reagent manufacturer’s certificate of analysis (https://www.thermofisher.com/order/catalog/product/90406) for lot number SE240163[93]. Proteins identified with 2 or more peptides were used for further statistical analysis. Comparison of protein levels in each strain were assessed for significances using a two-sample moderated T test was performed with an adjusted p-value threshold of less than 0.05 for assessing significantly altered proteins. For visualization purposes in Figure 2, the protein ratio associated with LepA was excluded from the volcano plot.

#### RNA Sequencing

Samples were sequenced on an Illumina HiSeq 2500 in paired-end mode with a read length of 125 bp. Approximately 4 million reads were collected for each sample. Reads were mapped to the genome sequence of *M. smegmatis* mc^2^ 155 as a reference genome using BWA (Li and Durbin, 2009). A python script was used to separate reads in .sam files that mapped to the positive strand and negative strand of the chromosome. Then reads mapping to each ORF (in a strand-specific manner) were tabulated. The raw read counts were converted to FPKMs (fragments per kilobase per million reads) by dividing by gene length (in bp) and total reads in the sample, and scaling up by 10^9. For analyses of differential gene expression, DESeq2 [94] was used to estimate log-fold-changes according to a hierarchical model based on the Negative Binomial distribution, and p-values were calculated via a Wald test as a measure of significance. P-values were adjusted for a false-discovery rate (FDR) of 5% over all genes by the Benjamini-Hochberg procedure. The raw sequence files are deposited in SRA under accession number SRP183056, and the gene expression levels (FPKMs) are deposited in GEO under accession number GSE126130.

### Luciferase assays

Strains were grown to log-phase and luciferase assays were conducted using Nanoglo Luciferase Assay System (Promega, Fitchburg, WI). Briefly, 100 μL of cells were mixed with 100 μL of Nanoglo reagent (prepared as kit protocol described). Within 2 minutes, luminescence measurements were taken in a TECAN Spark 10M plate reader with integration time of 1000ms. OD_600_ was also measured in each well, and luminescent values were normalized by OD_600_ to obtain relative luminescence values.

### Porin knockdown and contribution to LepA phenotype

Strains were grown to log-phase, diluted back into media with or without aTc, and allowed to grow for 15 hours to reach log-phase. Cells were stained with calcein-AM and analyzed by flow cytometry in biological triplicate, as described above.

### Ethidium bromide uptake assay

Strains were grown to log-phase, washed with PBS-0.4% glycerol (PBS-G) and prepared at an OD_600_ of 0.8. 100 μL of each strain was mixed with 100 μL 4 μg/mL of ethidium bromide (prepared in PBS-G) in a 96-well plate. Fluorescence was measured in a TECAN Spark 10M plate reader, using an excitation of 520 nm and emission of 600 nm.

### Fluorescence Microscopy and Image Analysis

Still imaging of MspA-mRFP strains was performed using a Nikon TI-E microscope at 60x magnification for image analysis and at 100x magnification to generate representative images. To quantify intensities, fluorescence intensities of single mycobacterial cells, a custom semi-automated ImageJ macro was run. As mycobacteria tend to clump, a user picked single cells from phase contrast images with the ImageJ ‘point tool’. Then, using ImageJ macros, a circular region of interest was created around each point, encompassing the bacterial cell and saved to the ROI manager. Automatic thresholding, and segmenting was performed to measure the intensity on a corresponding fluorescence image. This was repeated for each single-cell the user identified and the fluorescence intensities and areas were saved to text file for further analysis.

### Experimental Replicates

Unless otherwise noted all experiments were conducted at least twice, in biological triplicate.

### Data Analysis

Protein functional enrichment analysis was performed using ‘Functional Annotation’ software within the DAVID platform (https://david.ncifcrf.gov/home.jsp). Specifically, InterPro terms considered biologically significant, and therefore visualized, refers to at least 6 proteins in the list of 80 proteins significantly altered by LepA. Statistical significance was determined using a modified Fisher’s exact test[50]. Codon frequency was computed using custom R code, and specifically, the ‘seqinr’ package to analyze codon bias on a gene-by-gene and genome-wide basis. All other statistical measurements and tests are specified in the figure legends.

### mRNA stability assay

Cells were grown to mid-log phase in 30 mL, and aliquoted into 5 mL in 15 ml shaking Falcon tubes (3 replicates for each time point and each strain). At time zero, rifampin was added to replicate tubes at 200 μg/mL to inhibit transcription. Three replicates for each strain were flash frozen immediately. At 20 minutes, cells were again flash frozen. To harvest RNA, all cells were thawed on ice and processed as described above. mRNA levels at each time point for *mspA* and *sigA* were calculated using the ΔΔCt method, where each transcript was normalized by 16S rRNA levels [95].

### mRNA quantification

mRNA was quantified as described previously [60]. Briefly, purified RNA (DNase-treated) was used as template for cDNA synthesis, following manufacturer’s instructions with Superscript V (Life Technologies). RNA was removed from the reaction using alkaline hydrolysis and cDNA was cleaned using column purification (Zymo). qPCR was performed on purified cDNA using iTaq Universal SYBR Green Supermix (BioRad). mRNA fold change was calculated using the ΔΔCt method, where porin transcript level was normalized by *sigA* level in each genetic background.

## Supplemental Figure Legends

**Supplementary Figure 1**

a-b) Msm strains were treated with 10X concentrations of linezolid and isoniazid, and cell survival was measured by colony forming units per mL (CFU/ml) as in Figure 1. Values indicate mean survival with error bars indicating standard deviation of three biological replicates. c) SDS-PAGE gel displaying Ni-NTA column purification of Msm LepA. Lane 1: cell lysate; Lane 2: cell pellet; Lane 3: wash 1 (30 mM imidazole); Lane 4: wash 4; Lane 5: elution fraction 2; Lane 6: elution fraction 3; Lane 7: elution fraction 4; Lane 8: elution fraction 5; Lane 9: elution fraction 6; Lane 10: elution fraction 7; Lane 11: BenchMark Pre-stained ladder; Lane 12: elution fraction 3 + DTT. d) *In vitro* translation was carried out using Venus mRNA with increasing concentration of LepA. ‘-LepA’ represents reactions with no purified LepA added. Each concentration of LepA reflects the molar ratio of added LepA to the ribosome. Curves represent mean fluorescence with error bars indicating the range across two technical replicates.

**Supplementary Figure 2**

a) Fold-change in enrichment of InterPro functional annotation terms in proteins altered by loss of LepA relative to the total Msm proteome. *** P<0.001, ** P<0.001, *P<0.05, calculated using a modified Fisher’s exact test. b) A Clustal Omega alignment of mycobacterial porin family proteins in Msm. Black boxes indicate unique peptides that mapped to a mycobacterial porin. Decreasing hue corresponds to decreasing percent identity across the four sequences. c) Three sgRNAs with different PAM strengths [60] were chosen to achieve maximal knockdown of each porin. Black lines around a bar indicate selection of that sgRNA for future experiments. Fold repression, quantified using RT-qPCR, indicates the difference in knockdown between induced and uninduced strains. d) Visualization of MspA-mRFP in Msm. On the left, representative images of MspA-mRFP localization in cells with or without LepA. Scale bar represents 5 microns. On the right, quantification of average fluorescence across a single cell (N=100) from each strain. ***P<0.001 calculated using a Mann-Whitney test.

**Supplementary Figure 3**

a) For glycine codons, individual codon frequencies were calculated for all four porin genes, compared to genomic codon frequencies. b) The *mspA* gene was recoded at 8 glycine codons. Each glycine codon was selected based on the presence of a ‘GGT’ in *mspA-C* and a corresponding ‘GGC’ in *mspD*. Each number denotes the location of the T to C mutation made at the end of the glycine codon in reference to the *mspA* sequence. The glycine-recoded *mspA* was fused to a luciferase reporter. c) Luminescence of glycine-recoded *mspA* reporter compared to WT *mspA*. d) Luminescence of signal sequence reporters in *lepA*+ (filled bar) and *lepA*- (open bar) background. c-d) Values indicate mean of three replicates with error bars indicating standard deviation. e) ClustalW alignment of *mspA* (black letters) and *mspD* (dark gray letters) nucleotide sequences with points of fusion (black triangles) for chimeras in Figure 5D. Differences in *mspD* relative to *mspA* are highlighted in red.

## Supplementary Material

**Supplemental Dataset 1: Quantitative proteomics data**

Protein-level relative abundance of proteome in *lepA* strains

**Supplemental Dataset 2: RNA sequencing data**

DEseq output and RNA sequencing transcript abundance for *lepA* strains

**Supplementary Table 1: Drug susceptibility of *lepA* mutant**

Susceptibility of Msm strains was determined using a minimum inhibitory concentration (MIC) assay. Values are averages of three replicates, in μg/ml concentrations of drug.

**Supplemental Table 2: Strain List**

A description of bacterial strains used in the study

**Supplemental Table 3: Plasmid List**

A description of plasmids used in the study

**Supplemental Table 4: Primer List**

A description of primers used in the study

**Supplemental Table 5: Recombinant DNA**

A description of recombinant DNA used for this study

